# Primary cilia promote the differentiation of human neurons through the WNT signaling pathway

**DOI:** 10.1101/2023.04.06.535895

**Authors:** Andrea Coschiera, Masahito Yoshihara, Gilbert Lauter, Sini Ezer, Juha Kere, Peter Swoboda

**Affiliations:** Karolinska Institute, Department of Biosciences and Nutrition, Huddinge, Sweden; University of Helsinki, Stem Cells and Metabolism Research Program, and Folkhälsan Research Center, Helsinki, Finland; present address: Chiba University, Graduate School of Medicine, Department of Artificial Intelligence Medicine, Chuo-ku, Chiba-shi, Chiba, Japan; present address: Uppsala University, Department of Immunology, Genetics and Pathology, Rudbeck Laboratory, Uppsala, Sweden; equal contribution – joint first authorship: AC MY; equal contribution – joint last authorship: JK PS

## Abstract

Primary cilia emanate from most human cell types, including neurons. Cilia are important for communicating with the cell’s immediate environment: signal reception and transduction to/from the ciliated cell. Deregulation of ciliary signaling can lead to ciliopathies and certain neurodevelopmental disorders. In the developing brain cilia play well-documented roles for the expansion of the neural progenitor cell pool, while information about the roles of cilia during post-mitotic neuron differentiation and maturation is scarce. We employed ciliated Lund Human Mesencephalic (LUHMES) cells in time course experiments to assess the impact of ciliary signaling on neuron differentiation. By comparing ciliated and non-ciliated neuronal precursor cells and neurons in wild type and a mutant with altered cilia (RFX2 -/-), we discovered an early-differentiation “ciliary time window” during which transient cilia promote axon outgrowth, branching and arborization. Cilia promote neuron differentiation by tipping WNT signaling toward the non-canonical pathway, in turn activating WNT pathway output genes implicated in cyto-architectural changes. We provide a mechanistic entry point into when and how ciliary signaling coordinates, promotes and translates into anatomical changes. We hypothesize that ciliary alterations causing neuron differentiation defects may result in “mild” impairments of brain development, possibly underpinning certain aspects of neurodevelopmental disorders.

## INTRODUCTION

The primary cilium is an antenna-like structure projecting off polarized cell surfaces. It contains and elongates a microtubule-based axoneme from a modified mother centriole, the basal body, which is paired with a daughter centriole. The pair of centrioles is also part of the centrosome, a major cellular microtubule-organizing center (MTOC) important for mitosis (Joukov and De Nicolo, 2019), restricting ciliogenesis to the post-mitotic G1 (or G0) phase of the cell cycle and cilia disassembly to prior to the next mitosis (Ishikawa and Marshall, 2011).

Ciliogenesis is regulated by FOXJ1 and by members of the RFX transcription factor (TF) family (Senti and Swoboda, 2008; Piasecki et al., 2010; De Stasio et al., 2018), and mutations in these ciliogenic genes cause ciliary phenotypes (Choksi et al., 2014; Harris et al., 2021). In human there are eight RFX genes: RFX1-4 and RFX7 are widely expressed, including in brain tissues and the spinal cord (Sugiaman-Trapman et al., 2018); RFX2 is broadly required for ciliogenesis during vertebrate development (Chung et al., 2012).

Cilia harbor signal reception and transduction proteins. They act as cellular sensors, but also as senders of signals to the cell’s immediate environment (Garcia et al., 2018). Cilia coordinate and transduce a variety of essential signaling pathways (Wheway et al., 2018). Among these, WNT signaling plays a key role during embryogenesis and adult tissue homeostasis (Steinhart and Angers, 2018). Canonical and non-canonical WNT signaling is activated by distinct endogenous and exogenous ligands (Flores-Hernández et al., 2020). The main mediator of canonical WNT signaling is β-catenin, which accumulates in the cytoplasm and then translocates to the nucleus. There, β-catenin activates T-cell factor/lymphoid enhancer factor (TCF/LEF) TFs, in turn affecting target / WNT signaling output gene expression (MacDonald et al., 2009). Non-canonical, β-catenin independent WNT signaling activates the cytoplasmic β-catenin destruction complex, and thereby a different TF output complex called Activator protein 1 (AP-1) (Nishita et al., 2019; Arredondo et al., 2020). Through its interaction with the β-catenin destruction complex the ciliary protein Inversin modulates the balance of WNT pathways by tipping it toward non-canonical WNT signaling (Veland et al., 2013; Simons et al., 2015). A switch from canonical to non-canonical WNT signaling has been reported to mediate neural stem cell differentiation (Bengoa-Vergniory et al., 2014) and cilia may be the main cellular organelle initiating that switch.

Cilia are present on most human cell types, including neurons, and defective cilia can lead to conditions known as ciliopathies, which display pleiotropic clinical, including brain, phenotypes (Suciu and Caspary, 2021; Tereshko et al., 2022). Also, certain neurodevelopmental conditions and disorders such as dyslexia (Tammimies et al., 2016; Bieder et al., 2020), autism and schizophrenia (Reiter and Leroux, 2017; Lauter et al., 2018), display strong connections to ciliary aberrations, including through disease-associated candidate genes with demonstrated ciliary functions.

During embryogenesis, cilia extend apically from the neuroepithelium into the lumen of the neural tube to detect morphogens gradients (Lepanto et al., 2016). Cilia direct the transition of neuroepithelial stem cells into neural progenitor cells (neurogenesis) that will give rise to all neuronal and glial cell types of the central nervous system (Liu et al., 2021). Cilia as sensory and signaling devices have well-documented roles in the initial expansion of the neural progenitor cell pool (Liu et al., 2021; Suciu and Caspary, 2021), and in neuron migration (Matsumoto et al., 2019; Stoufflet et al., 2020). Also, recent reports demonstrated that cilia impact axon navigation (SHH signaling; Toro-Tapia and Das, 2020), growth cone and axon tract development (PI3K/AKT signal transduction; Guo et al., 2019), the shaping of dendritic structures (AC3 signal transduction; Guadiana et al., 2013), and interneuronal connectivity (GPCR signaling; Guo et al., 2017).

In parts concurrent with neuron migration, neuron differentiation follows a pattern of successive stages, encompassing precursor cell polarization, axonogenesis, axon outgrowth, elongation and branching, the formation of dendrites, spines and synapses, including pruning steps, and finally the formation of neuronal circuits (Kaech et al., 2012; Meka et al., 2020; Meka et al., 2022). The impact of cilia on (early) neuron differentiation is still under debate, while centrosome and Golgi apparatus have already been shown to be crucial for microtubule nucleation during polarization after mitosis and for axon specification and axonogenesis (Meka et al., 2020; Meka et al., 2022).

Here we investigate cilia in a human neuronal cell line, LUHMES (Lauter et al., 2020), where cells proliferate as neuronal precursors or can be induced to differentiate and mature into neurons. Upon induction to differentiate, ciliation occurs asynchronously, is transient during the early differentiation stages and not every neuron ciliates. We determined in individual neurons how and when cilia impact neuron differentiation; by comparing ciliated and non-ciliated neurons in wild type (WT) and in an RFX2 -/- mutant with ciliary alterations. We discovered a “ciliary time window” during which cilia promote axon outgrowth, branching and arborization. Cilia promote these neuron differentiation steps by tipping WNT signaling toward the non-canonical pathway, in turn activating WNT pathway output genes implicated in cyto-architectural changes. Thereby we provide a mechanistic entry point into how and when during neuron differentiation ciliary signaling coordinates, promotes and translates into anatomical changes. The emerging, appropriate neuron anatomy is a prerequisite for establishing functional connections (circuit formation). Our findings suggest a critical spatiotemporal role of cilia in neuron and brain development. We hypothesize that ciliary alterations causing neuron differentiation defects may result in “mild” impairments of brain development, possibly underpinning the onset of certain aspects of neurodevelopmental disorders.

## RESULTS

### Cilia promote neuron differentiation: axon outgrowth

In the LUHMES cell model (Lauter et al., 2020), we used unsynchronized neuronal precursor cell proliferation conditions and timed (d0-d1), change of culture medium-induced release into neuron differentiation. We noticed that ciliated neurons more efficiently accomplished axon outgrowth as compared to non-ciliated neurons.

To quantify these differentiation differences, we performed stringent time course experiments, where we assessed established neuron differentiation and outgrowth milestones: stage 1 (growth cone protrusions), stages 2a and 2b (bipolar stage, engorgement and consolidation), stage 3 (axon outgrowth, break of symmetry), and stages 4 and 5 (maturation, formation of dendrites, spines and synapses) (Meka et al., 2020; Meka et al., 2022) (Figure 1A). LUHMES neurons differentiate through these stages within about one week (Lauter et al., 2020) (Figure 1B-F). We performed immunocytochemistry using markers for cilia (ARL13B), centrosomes/basal bodies (PCNT) and for axons (TAU, TRIM46) (Larkins et al., 2011; Mühlhans et al., 2011; Bell et al., 2021) to quantify the presence of cilia and emerging axons (stage 3) throughout the entire LUHMES neuron differentiation protocol (d0-d6).

**Figure 1:**
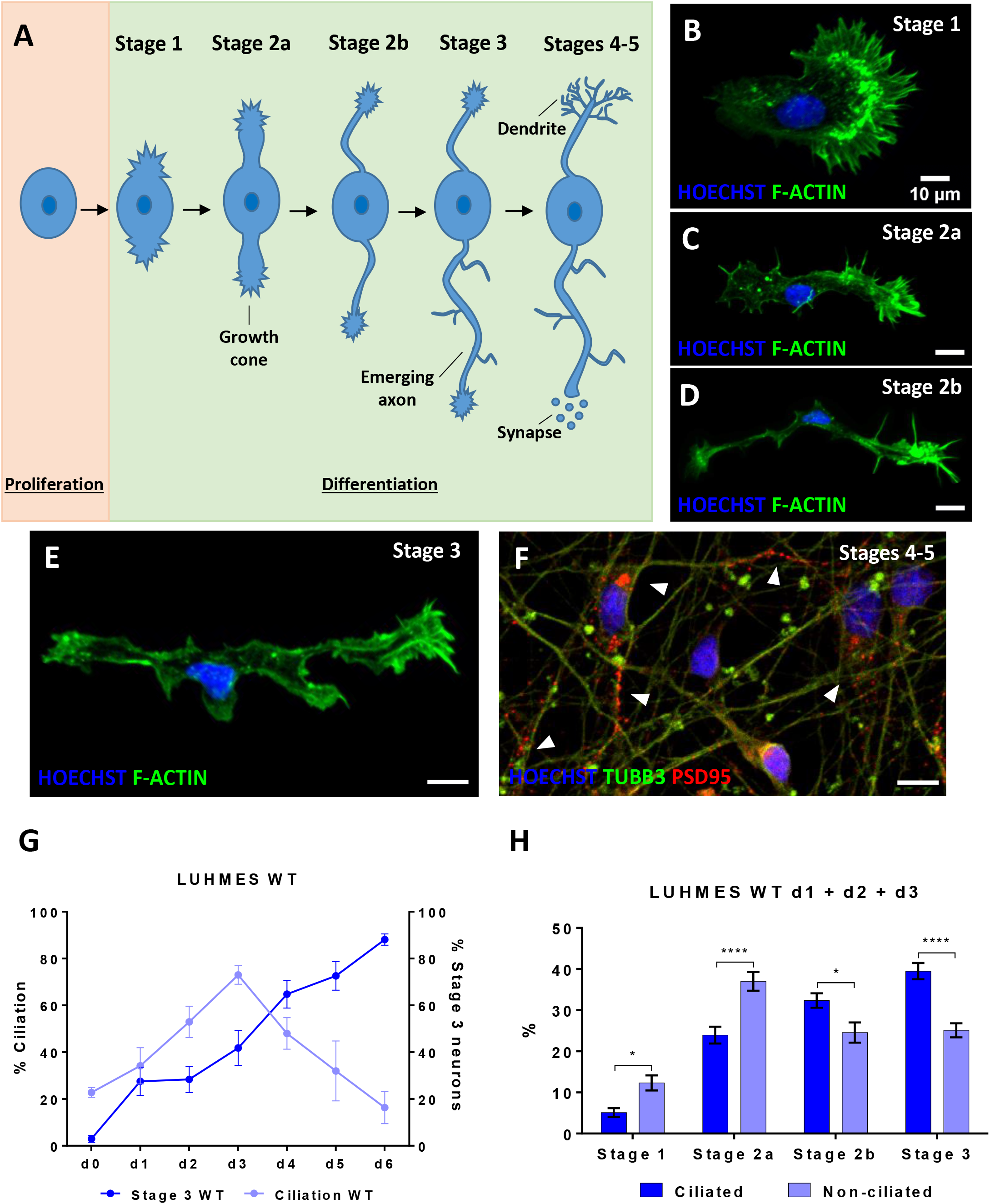
Stages of neuron differentiation: cilia promote axon outgrowth. See also Supplementary Figures S1 and S2. **(A)** Schematic illustration of neuron differentiation stages (Meka et al., 2020): stage 1 – growth cone protrusions during neuron polarization; stage 2a – symmetric bipolar stage with microtubule invasion into growth cones, first neurite extension; stage 2b – symmetric bipolar stage with microtubule bundles consolidation, further neurite extension; stage 3 – the emerging axon outgrows the opposite neurite breaking the bipolar symmetry; stages 4 and 5 – dendrites and dendritic spines formation, further maturation with formation of synapses. **(B-F)** Staging of human LUHMES neuron differentiation: immunocytochemistry detects nuclei (Hoechst staining), the cytoskeleton (Phalloidin stains F-actin), microtubule bundles (TUBB3) and emerging synapses (PSD95). **(G)** The percentage of ciliation in differentiating human LUHMES neurons increases once neurons exit the cell cycle (d1), peaks in differentiating neurons (d3) and then decreases during the later stages of neuron differentiation and maturation (d4-d6). The percentage of neurons reaching stage 3 of differentiation steadily increases throughout the entire differentiation and maturation process (n=134-394). **(H)** Populations (d1-d3) of ciliated LUHMES neurons (n=738) are more efficient than non-ciliated LUHMES neurons (n=573) in breaking the bipolar symmetry of stages 2a and 2b, and thus, in promoting axon outgrowth. Mean values ± s.e.m. are shown. The results are from a minimum of three independent experiments with a total of at least six technical replicates. We conducted regular two-way ANOVA analyses (not repeated measures) with multiple comparisons (Bonferroni’s test) between groups. *p<0.05; ****p<0.0001.

We released LUHMES precursor cells into neuron differentiation from an unsynchronized cell cycle starting point (d0/d1). Thereby some cells exit the cell cycle, become post-mitotic and ciliate earlier than others during the initial phases of differentiation. Ciliation increased steadily during early differentiation, peaking at d3 with most neurons (about 70%) having ciliated. Detectable ciliation then decreased during the later phases of differentiation (d4-d6) (Figure 1G). For each differentiation time point (d0-d6), we also determined the number of neurons in the population that were able to break the bipolar symmetry and initiate the outgrowth of the future axon (stages 2 and 3). The percentage of neurons with a defined, emerging axon steadily increased over time, consistent with the ongoing maturation of neurons during the differentiation process (Figure 1G, Supplementary Figure S1). At a population level, ciliation “precedes” axonogenesis and axon outgrowth (Figure 1G).

In these populations of differentiating LUHMES neurons we then compared ciliated and non-ciliated neurons with regard to how efficiently they reach differentiation stage 3 during the initial phases of the differentiation process (d1-d3). We found that the majority of neurons that reach differentiation stages 2b and 3 were neurons with a cilium, while for the earlier differentiation stages 1 and 2a the non-ciliated neuron subpopulation was in the majority, at all three time points examined (d1-d3) ( Figure 1H, Supplementary Figure S2A-D).

Taken together our results indicate that the presence of cilia and ciliary signaling promote neuron differentiation and maturation during the initial phases, constituting a “transient ciliary time window”, exemplified by a crucial feature, break of symmetry and axon outgrowth.

### Cilia structure is altered in human LUHMES RFX2 -/- (knockout) cells and neurons

To experimentally manipulate differences in differentiation between ciliated and non-ciliated neurons, we altered cilia by mutation. We knocked out the gene for the ciliogenic transcription factor (TF) RFX2. The RFX family of TFs is essential for ciliogenesis (Choksi et al., 2014). In vertebrates *RFX2* gene function is required for ciliogenesis during development (Chung et al., 2012), in humans *RFX2* is abundantly expressed in the brain (Sugiaman-Trapman et al., 2018), and in human LUHMES neurons *RFX2* is highly expressed during early differentiation (d0-d2/d3) (Lauter et al., 2020).

We used CRISPR/Cas9 with guide RNAs (gRNAs) specific for sequences upstream of the exons encoding the RFX2 DNA binding domain (DBD), essential for protein (TF) function. Thereby we generated LUHMES RFX2 -/- deletion alleles that lead to translational frame shifts and premature stop codons, destroying the DBD and all subsequent protein domains, thereby constituting a gene knockout mutation (Figure 2A, Supplementary Figure S3A-D). We demonstrated by qRT-PCR that *RFX2* mRNA expression is strongly downregulated (Supplementary Figure S3B), and by Western blot that RFX2 protein is absent (Figure 2B), throughout the entire neuron differentiation time course.

**Figure 2:**
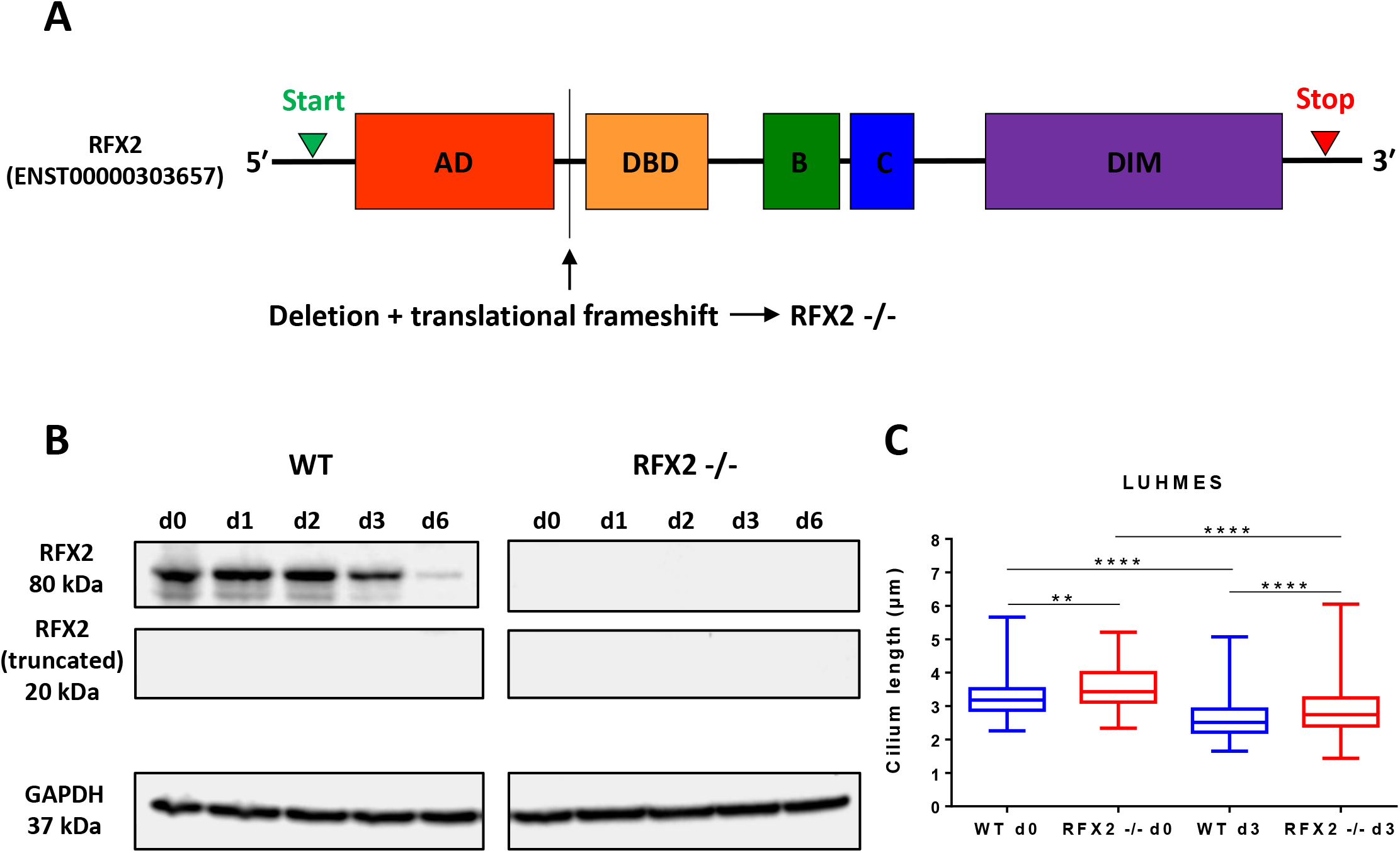
Cilia structure is altered (cilia are longer) in human LUHMES RFX2 -/- mutants. See also Supplementary Figure S3. **(A)** Schematic illustration of the RFX2 transcription factor (TF) gene structure with functional protein domains indicated: AD – activation domain, DBD – DNA binding domain, B and C – domains B and C of unknown function, DIM – dimerization domain. CRISPR/Cas9-engineered, small deletions lead to reading frame shifts upstream of the RFX2 DBD, essential for TF functionality, thereby creating an RFX2 -/- knockout mutant. **(B)** Western blot analyses of RFX2 protein expression in LUHMES wild-type (WT) and RFX2 -/- mutants: In the RFX2 -/- mutant RFX2 protein expression is completely absent during early neuron differentiation time points (d1-d3) and at the end of differentiation and maturation (d6). Also, no protein expression of a hypothetical, predicted truncated RFX2 protein product was detectable as a result of CRISPR/Cas9 mutagenesis. The housekeeping protein GAPDH was used as a loading control. **(C)** Cilia of RFX2 -/- mutants are longer than WT cilia, both at the neuronal precursor cell stage (d0) and in differentiating neurons (d3). Immunofluorescence-based length measurements: ARL13B marks the ciliary shaft, PCNT marks the basal body. Statistics: WT d0 (n=74), RFX2 -/- d0 (n=119), WT d3 (n=210), RFX2 -/- d3 (n=218). Data are shown in Box and Whisker plots (min to max) from at least three independent experiments. We conducted regular one-way ANOVA analyses with multiple comparisons (Bonferroni’s test) between groups. **p<0.005; ****p<0.0001.

In control experiments we then evaluated whether the RFX2 -/- genotype caused any overt growth or gross anatomical abnormalities in differentiating neurons. We found this not to be the case. Both WT and RFX2 -/- neurons displayed near-identical growth rates, whether grown in regular growth medium for the proliferation of the precursor cell stage or when grown in growth and differentiation medium for the induction of neuron differentiation (Supplementary Figure S3E-F). Neither did we find any relevant differences between WT and RFX2 -/- neurons in overall neuron anatomy during the early phases of differentiation (d1-d3). Using whole-cell immunocytochemistry (neurites were marked with either anti-TAU or anti-TRIM46) we detected that both cultures were dominated to near-identical extents by bipolar, followed by multipolar and then unipolar neurons (Supplementary Figure S3G-H).

To determine the impact of the RFX2 -/- mutations on ciliary structure we used confocal microscopy and as a first measure determined ciliary length, which we found to be significantly longer than in WT (Figure 2C). Taken together, our observations, expectedly, point toward a ciliary role of RFX2, while the cell cycle and cell proliferation, and gross neuron anatomy appear to be unaffected by mutation of RFX2.

### Absent or altered cilia affect several aspects of human LUHMES neuron differentiation

Our experimental setup allowed for direct comparisons between differentiating neurons: ciliated *versus* altered cilia *versus* non-ciliated. Using immunocytochemistry, we focused mostly on axon-associated phenotypes during the entire time course of neuron differentiation and maturation (d0-d6).

Using a marker for cilia (ARL13B) we observed that ciliation was similarly transient for both WT and RFX2 -/- (Figure 3A). Using markers for axons (TAU, TRIM46) we found consistently that a population of WT neurons more efficiently reached differentiation stage 3 as compared to a population of RFX2 -/- neurons (Figure 3A). It appears that (functional) WT cilia provide a differentiation “advantage” over altered (RFX2 -/-) or absent cilia to facilitate axon outgrowth (Figure 3A, Supplementary Figure S2). Using Phalloidin staining to detect cytoskeletal F-actin we then investigated aspects of emerging axon tract development such as axon branching and arborization in both ciliated and non-ciliated differentiation stage 3 neurons. We found in WT ciliated neurons the emerging axons to be significantly more branched than in non-ciliated neurons (Figures 3B, 3D). These differences in axon branching and arborization were absent in RFX2 -/- when comparing neurons with altered and neurons with absent cilia (Figure 3C).

**Figure 3:**
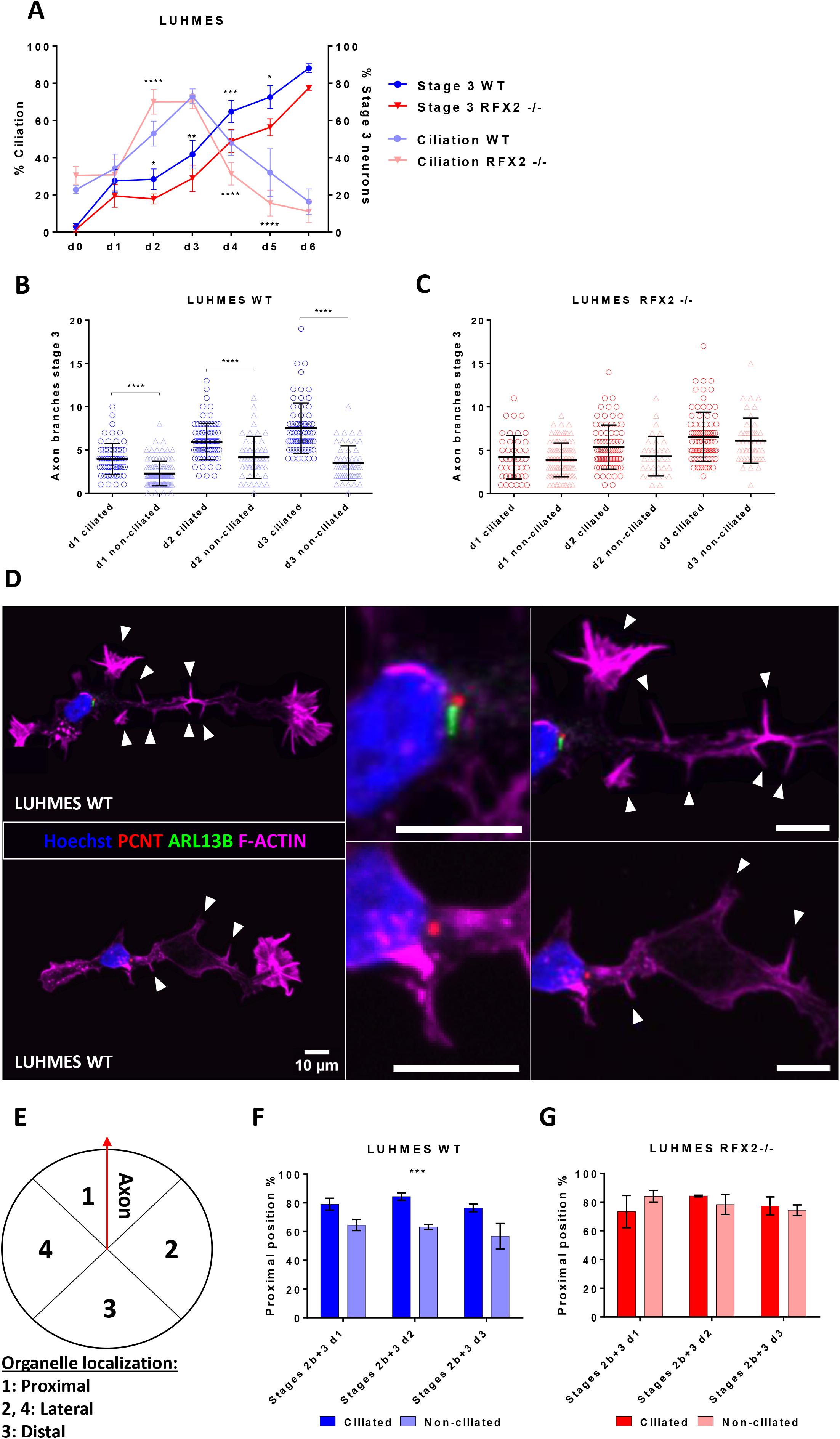
Altered cilia affect different aspects of human LUHMES neuron differentiation: axon outgrowth and branching pattern, and the spatial relationship between cilium and axon. See also Supplementary Figures S2 and S4. **(A)** The percentage of ciliation in differentiating human LUHMES neurons increases and declines slightly earlier in RFX2 -/- mutants as compared to WT neurons. The percentage of neurons reaching stage 3 of differentiation is delayed in RFX2 -/- mutants as compared to WT neurons throughout the entire differentiation and maturation process (WT n=134-394; RFX2 -/- n=162-492). **(B-C)** Axons of ciliated neurons are more branched and arborized as compared to non-ciliated neurons in WT, whereas in RFX2 -/- mutants there is no difference between ciliated and non-ciliated neurons. **(D)** Detection by immunocytochemistry demonstrates the effect the cilium imparts on the complexity of axon branching in WT neurons: top – ciliated neuron, bottom – non-ciliated neuron (arrowheads indicate axon branch points); Hoechst stain – nucleus, PCNT – centriole/basal body marker, ARL13B – ciliary marker, Phalloidin – cytoskeleton/F-actin marker. **(E)** Primary cilia are anchored at the cell surface by a basal body, a modified centriole. Schematic illustration of the neuronal cell body: possible cellular localization of cilia and centriole/basal body (ciliated neuron) or of centriole-only (non-ciliated neuron), in relation to the emerging axon outgrowth position (red line). **(F-G)** In WT, the typical axon-proximal localization of centrioles (centrosome) is observed more frequently (in %) in ciliated neurons than in non-ciliated neurons (ciliated n=446; non-ciliated n=170). In RFX2 -/- mutants this difference (in %) between ciliated and non-ciliated neurons disappears (ciliated n=214; non-ciliated n=169). Mean values are shown ± s.d. **(A-C)** and ± s.e.m **(F-G)**. The results are from a minimum of three independent experiments with a minimum of two technical replicates each. We conducted regular two-way ANOVA analyses (not repeated measures) with multiple comparisons (Bonferroni’s test) between groups. *p<0.05; **p<0.005; ***p<0.0005; ****p<0.0001.

Primary cilia emanate from the basal body, a modified mother centriole, which together with the daughter centriole forms the centrosome. The centrioles of the centrosome have been shown to play a key role in the initial steps of neuron differentiation by determining polarization and axonogenesis (Meka et al., 2020; Meka et al., 2022). Comparing ciliated and non-ciliated neurons in WT and RFX2 -/- allows for separating the different contributions of cilia and centrioles during the time course of neuron differentiation. Immunocytochemistry of stage 2b and stage 3 neurons (cf. Figure 1A) with a marker for centrosomes/centrioles/basal bodies (PCNT) and a marker for cilia (ARL13B) revealed that the alignment of centriole localization with the axis of axon outgrowth (Figure 3E) was, in WT, significantly stronger in ciliated neurons as compared to non-ciliated neurons (Figure 3F, Supplementary Figure S4). Again, these differences between ciliated and non-ciliated neurons disappeared in RFX2 -/- (Figure 3G, Supplementary Figure S4).

We conclude that the presence of functional primary cilia promotes several aspects of neuron differentiation: the precision of centriole localization in connection to the axis of axon outgrowth, axon outgrowth itself, and axon branching. While mutation of the ciliogenic RFX2 TF gene, altering cilia structure and function, delays and deregulates these aspects of the neuron differentiation process.

### RFX2 -/- knockout leads to delayed neuron differentiation also at the transcriptome level

To monitor gene expression during neuron differentiation, we subjected WT and RFX2 -/- LUHMES cells and neurons to modified single-cell tagged reverse transcription RNA-sequencing (STRT RNA-Seq), which captures the 5’ ends of transcripts (transcript far 5’ ends, TFEs) (Islam et al., 2011; Islam et al., 2014; Ezer et al., 2021). Cells were collected from d0 to d6 with 4–6 replicates per time point (Supplementary Table 1). Principal component analysis (PCA) revealed that RFX2 -/- neurons showed a delayed differentiation pattern as compared to WT. For example, the gene expression profile of d4 differentiating RFX2 -/- neurons displayed high similarity with d3 differentiating WT neurons (Figure 4A). For confirmation, we compared our STRT RNA-seq results with the single-cell RNA-seq data set from human fetal midbrain *in vivo* (La Manno et al., 2016). As shown in a previous study (Lauter et al., 2020), during the time course of differentiation (d0-d6) WT LUHMES neurons progressively acquired high similarity with differentiated human midbrain neurons. RFX2 -/- neurons, on the other hand, showed a pronouncedly weaker similarity with differentiating human midbrain neurons as compared to WT, and again, typically were a day “behind” in differentiation (Figure 4B).

**Figure 4:**
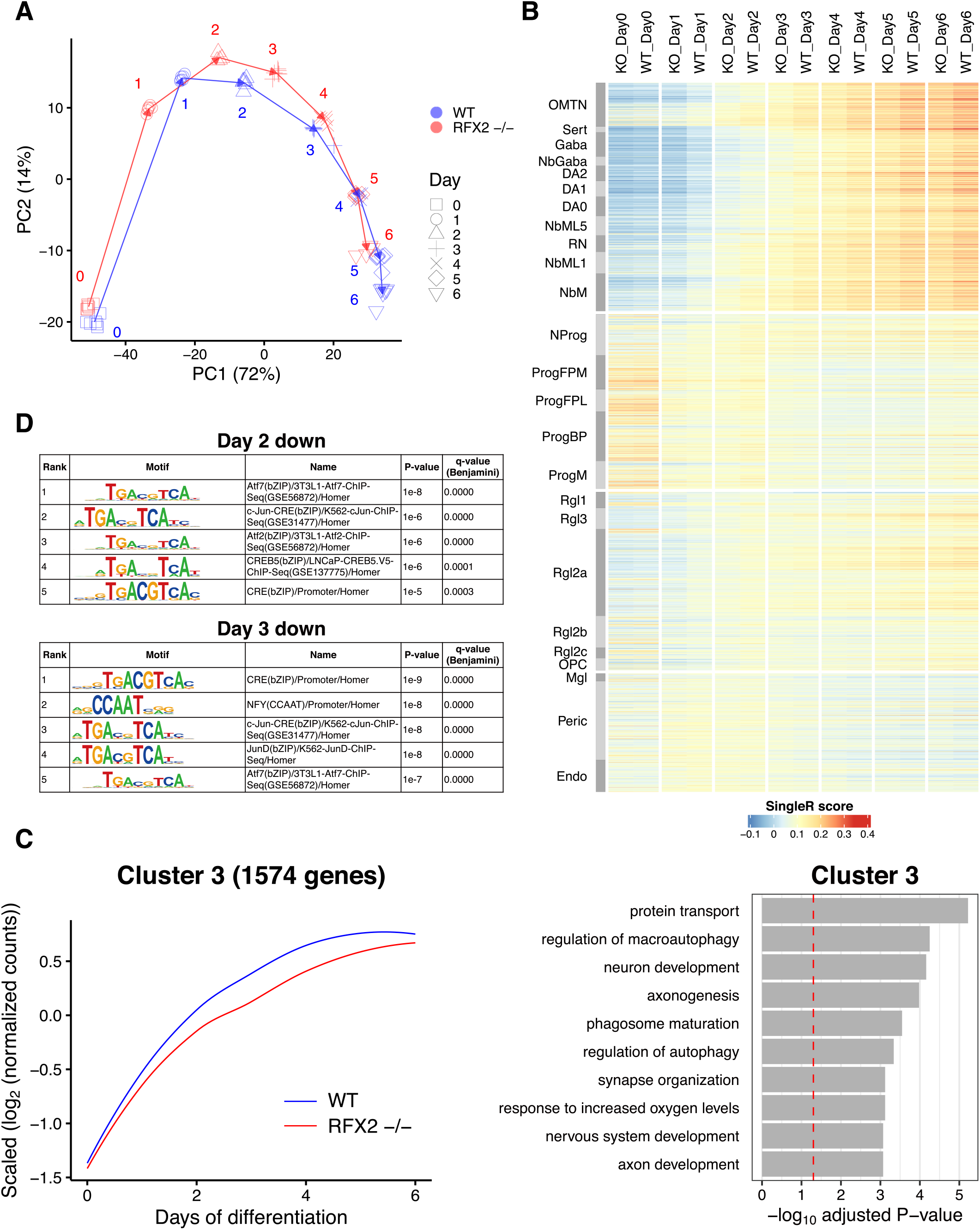
Transcriptomics confirm the delay in differentiation of human RFX2 -/- LUHMES neurons. See also Supplementary Figures S5-S8. **(A)** Principal component analysis of 78 (41 WT and 37 RFX2 -/-) LUHMES STRT RNA-seq time course samples, representing the process of neuron differentiation from d0 to d6. Numbers represent the days and arrows connect the neighboring days. **(B)** Similarity heat map of SingleR annotation scores of different types of human fetal midbrain cells and neurons using the 78 LUHMES WT and RFX2 -/- samples as reference. OMTN, oculomotor and trochlear nucleus; Sert, serotonergic neurons; Gaba, GABAergic neurons; NbGaba, GABAergic neuroblasts; DA, dopaminergic neurons; NbML, mediolateral neuroblasts; RN, red nucleus; NbM, medial neuroblast; NProg, neuronal progenitor; Prog, progenitor medial floor plate (FPM), lateral floor plate (FPL), basal plate (BP), midline (M); Rgl, radial glia-like cells; OPC, oligodendrocyte precursor cells; Mgl, microglia; Peric, pericytes; Endo, endothelial cells. The order of these clusters follows the order originally described in La Manno et al. (2016). Note the apparent differentiation delay of RFX2 -/-, visible in the comparisons to differentiating neuronal cell types (d1/d2-d6; upper right corner), while in the comparisons to precursor cell types (d0; middle left) no differences between WT and RFX2 -/- are apparent. **(C)** Left: Scaled expression profile of 1,574 genes downregulated in RFX2 -/- LUHMES STRT RNA-seq samples (Cluster 3) throughout the differentiation time course. 3,476 genes with a significantly variable expression between WT and RFX2 -/- throughout the time course were clustered into three clusters based on their expression patterns during the differentiation process. Lines are local polynomial regression fittings of the scaled expression of the genes in each cluster, depicted in blue (WT) and in red (RFX2 - /-). Right: Gene ontology (GO) enrichment analysis of these 1,574 genes in Cluster 3. The red dashed line represents the adjusted p-value = 0.05. Clusters 1-3 are depicted side-by-side in Supplementary Figure S5. **(D)** Transcription factor binding motif enrichment analysis of significantly downregulated transcript far 5’ ends (TFEs) in RFX2 -/- LUHMES STRT RNA-seq samples at day 2 and day 3: only the top five significantly enriched motifs are listed. The complete transcription factor binding motif enrichment analysis throughout the time course of neuron differentiation and maturation (d0-d6) is shown in Supplementary Figure S8.

To identify the temporal gene expression pattern between WT and RFX2 -/-, we clustered 3,476 significantly variable genes into 3 clusters based on the expression pattern over the time course (Figure 4C, Supplementary Figure S5). Genes in Cluster 3 were upregulated during neuron differentiation but lowly expressed in RFX2 -/- as compared to WT throughout the entire time course. Of note, these 1,574 genes in Cluster 3 were enriched with genes related with neuron development and axonogenesis (Figure 4C). *DCX* and *MAPT* (neuronal marker genes), *PAX6*, *TH*, and *SST* (neuronal function-related genes) were classified into Cluster 3 (Supplementary Figure S6A, S6B, Supplementary Table S2, S3). The gene ontology (GO) term ‘cilium assembly’ was also significantly enriched in this cluster (adjusted *P*-value = 0.005) (Supplementary Table S3). Prominent ciliary genes such as *IFT27*, *TUBB4A*, and *SEPT3* were classified into this cluster (Supplementary Figure S6C, Supplementary Table S2, S3). Further, we found that significantly downregulated genes in RFX2 -/- at each time point were enriched for the term ‘cilium assembly’ and ciliary genes, especially at d3 and d4 of neuron differentiation (Supplementary Figure S7, Supplementary Table S4). Together, these observations strongly support the importance of RFX2 for neuron development and ciliogenesis.

The STRT RNA-seq expression level of the *RFX2* gene was comparable between WT and RFX2 -/- (Supplementary Figure S6D), which should be the case, because the CRISPR/Cas9 gene editing we performed, did not affect the 5’ end of *RFX2* gene (Figure 2, Supplementary Figure S3A-D). To confirm reduced (or absent) RFX2 TF function in the RFX2 -/- background, we examined the enrichment of TF binding site motifs closely associated with significantly downregulated TFEs at each time point. RFX2 binding motifs (X-boxes) were most significantly enriched at d4 and d5 (Supplementary Figure S8), indicating that RFX2 target genes were downregulated in the RFX2 -/- background. We further discovered that binding sites for the Atf, Jun, and Fos families of TFs were significantly enriched at d2 and d3 (Figure 4D). These TFs are members of the AP-1 transcriptional complex that is activated by WNT signaling (Hankey et al., 2018; Liu et al., 2021). In turn, this suggests that RFX2 contributes to the activation of the WNT signaling pathway.

### WNT signaling is essential to promote neuron differentiation

Primary cilia act as transducers of signaling pathways important for cortex development and axon pathfinding (Hasenpusch-Theil et al., 2020), neuron migration (Stoufflet et al., 2020) and neuronal circuitry formation (Guo et al., 2019). Thus, malfunction of ciliary signaling may lead to ciliopathies with brain phenotypes, to neurodevelopmental disorders or brain conditions (Reiter and Leroux, 2017; Lauter et al., 2018; Ma et al., 2022). Neural and neuron differentiation results from crosstalk of multiple (ciliary) signaling pathways (Wheway et al., 2018), where WNT signaling was shown to play a highly relevant role (Inestrosa and Varela-Nallar, 2015), for example in neural stem cell differentiation (Bengoa-Vergniory et al., 2014). Our STRT RNA-seq data revealed LUHMES RFX2 -/- neurons to be delayed in differentiation, whereby a number of downregulated genes in RFX2 -/- appeared to be targets of TF families (Atf, Jun, Fos) that are involved in non-canonical WNT signaling (Figure 4D).

To determine whether ciliary WNT signaling has an impact on LUHMES neuron differentiation, we used a pathway inhibitor (Wnt-C59) (Motono et al., 2016) and examined its impact on the two main phenotypes we previously uncovered in differentiating neurons (at d3): axon outgrowth (how efficiently neurons reach differentiation stage 3) and axon branching/arborization. In WT the axon marker TAU highlighted a significantly reduced number of stage 3 neurons upon treatment with Wnt-C59 (Figure 5A), while no differences between control and Wnt-C59 treatment were observed in the RFX2 -/- background (Figure 5B). When comparing ciliated to non-ciliated neurons in WT, we revealed a strong ciliary contribution to promoting axon outgrowth (Figure 5C), which was abolished when WNT signaling was inhibited (Figure 5C) or under any condition in RFX2 -/- neurons (Figure 5D). We then used Phalloidin staining cytoskeletal F-actin to detect the shapes of entire neurons, including the number of axon branches/arborization. Again, WT ciliated neurons had more axon branches than non-ciliated neurons (Figure 5E), a difference that was abolished upon treatment with Wnt-C59, curiously though at a higher level of branching (Figure 5E), or under any condition in RFX2 -/- neurons (Figure 5F).

**Figure 5:**
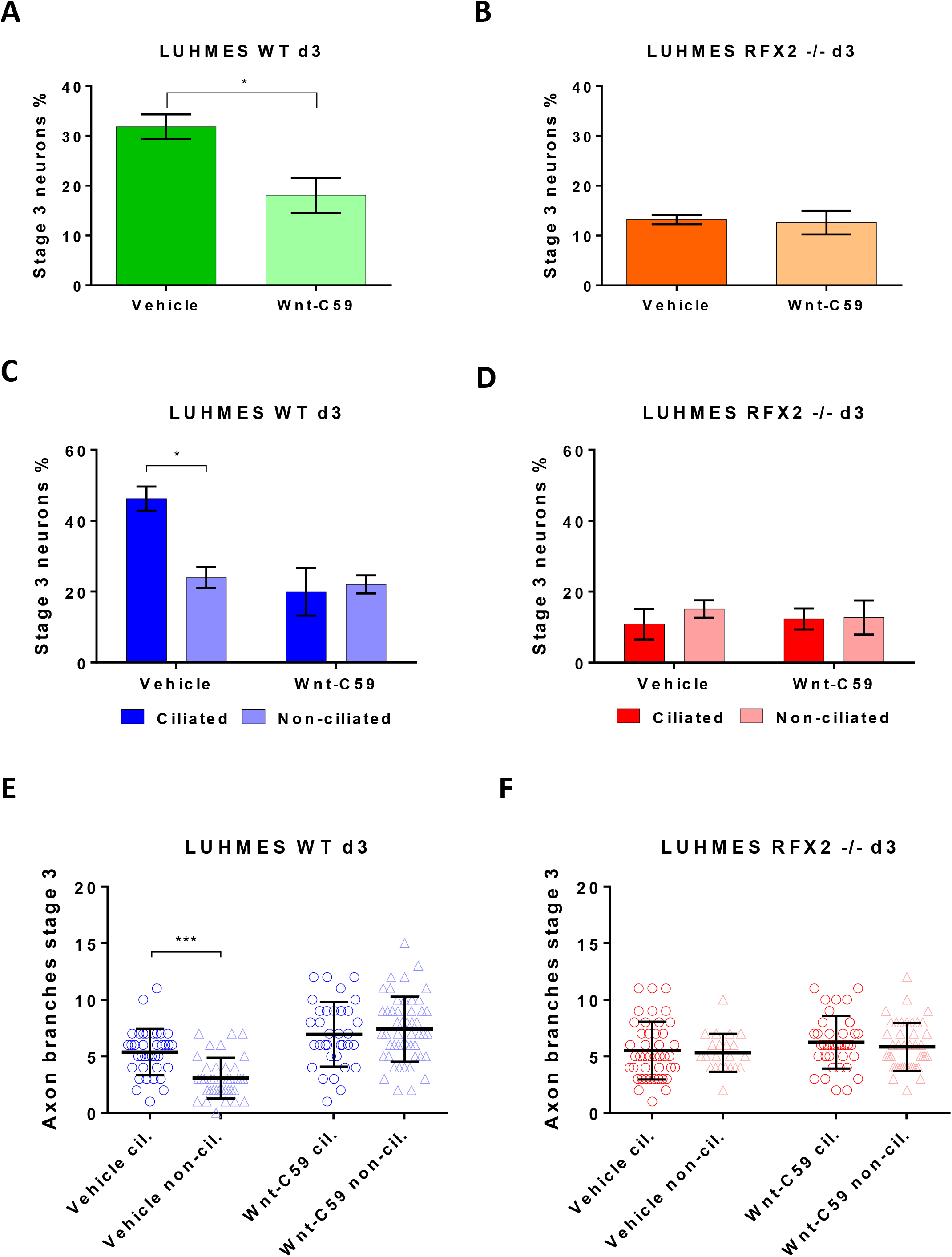
WNT signaling is essential for promoting human LUHMES neuron differentiation. **(A-B)** Inhibiting WNT signaling by treating LUHMES neurons with the antagonist Wnt-C59 negatively affects axon outgrowth (how efficiently neurons reach differentiation stage 3) in **(A)** WT but not in **(B)** an RFX2 -/- background. Control (vehicle): the same treatment, but without antagonist. **(C-D)** WT ciliated neurons reach differentiation stage 3 more efficiently than non-ciliated neurons in the control experiment (vehicle), while no difference was observed between ciliated and non-ciliated neurons after treatment with the antagonist Wnt-C59 **(C)**. No differences were observed in RFX2 -/- neurons **(D)** under the same experimental conditions (ciliated versus non-ciliated; with versus without antagonist Wnt-C59). **(E-F)** Wnt-C59 treatment deregulates axon branching. In the control experiment (vehicle), axon branching is promoted in WT ciliated neurons as compared to non-ciliated neurons. Upon treatment with the antagonist Wnt-C59 this difference is alleviated **(E)**. In RFX2 -/- neurons with altered cilia there are no differences in axon branching between ciliated and non-ciliated neurons in the control experiment (vehicle); likewise, for the treatment with the antagonist Wnt-C59 **(F)**. Mean values are shown ± s.e.m **(A-D)** and ± s.d. **(E-F)**. The results are from two independent experiments with a minimum of three technical replicates each. We conducted an unpaired two-sided t-test (95% confidence level) **(A-B)** and regular two-way ANOVA analyses (not repeated measures) with multiple comparisons (Bonferroni’s test) between conditions **(C-F)**. *p<0.05; ***p<0.0005.

These results indicate a direct involvement of WNT signaling in aspects of neuron differentiation, like axon outgrowth and branching. Disrupting WNT signaling by Wnt-C59 caused axon outgrowth and branching phenotypes that mimicked the outgrowth and branching phenotypes observed in control (vehicle) RFX2 -/- neurons with altered cilia, further strengthening the connection between WNT signaling and primary cilia for promoting neuron differentiation.

### Primary cilia regulate the subcellular localization of β-catenin, the main mediator of WNT signaling

How might primary cilia be involved in transducing WNT signaling? β-catenin is the main mediator for the activation of canonical WNT signaling and it is well expressed in both LUHMES WT and RFX2 -/- neurons (Supplementary Figure S6D). β-catenin can be active or inactive: the phosphorylated form is degraded by a multiprotein destruction complex in the cytoplasm; the non-phosphorylated, active form is stabilized and translocates into the nucleus to affect canonical WNT target genes expression (Shang et al., 2017). Cilia have been described as crucial regulators of the subcellular localization, and therefore activity, of β-catenin (Oh and Katsanis, 2013; May-Simera et al., 2018).

To determine how the cilia of LUHMES neurons influence β-catenin turnover and consequently WNT signaling, we quantified the expression of the active (non-phospho) form of β-catenin by Western blot. Given the ciliation pattern of LUHMES neurons during the early neuron differentiation time window (cf. Figure 1G), we analyzed active β-catenin during d1-d3 in both WT and RFX2 -/- backgrounds. Different from the overall β-catenin expression (Supplementary Figure S6D), we found a significant upregulation of the active form in RFX2 -/- on d2 and d3 (Figure 6A, 6B). Differentiating neurons at d1 did not result in significant differences between WT and RFX2 -/-, which may be due to the still low percentage of ciliation (about 30%) right after release from precursor cell proliferation into neuron differentiation, as compared to the following differentiation days (about 50% ciliation for d2 and 70% for d3) (cf. Figure 1G).

**Figure 6:**
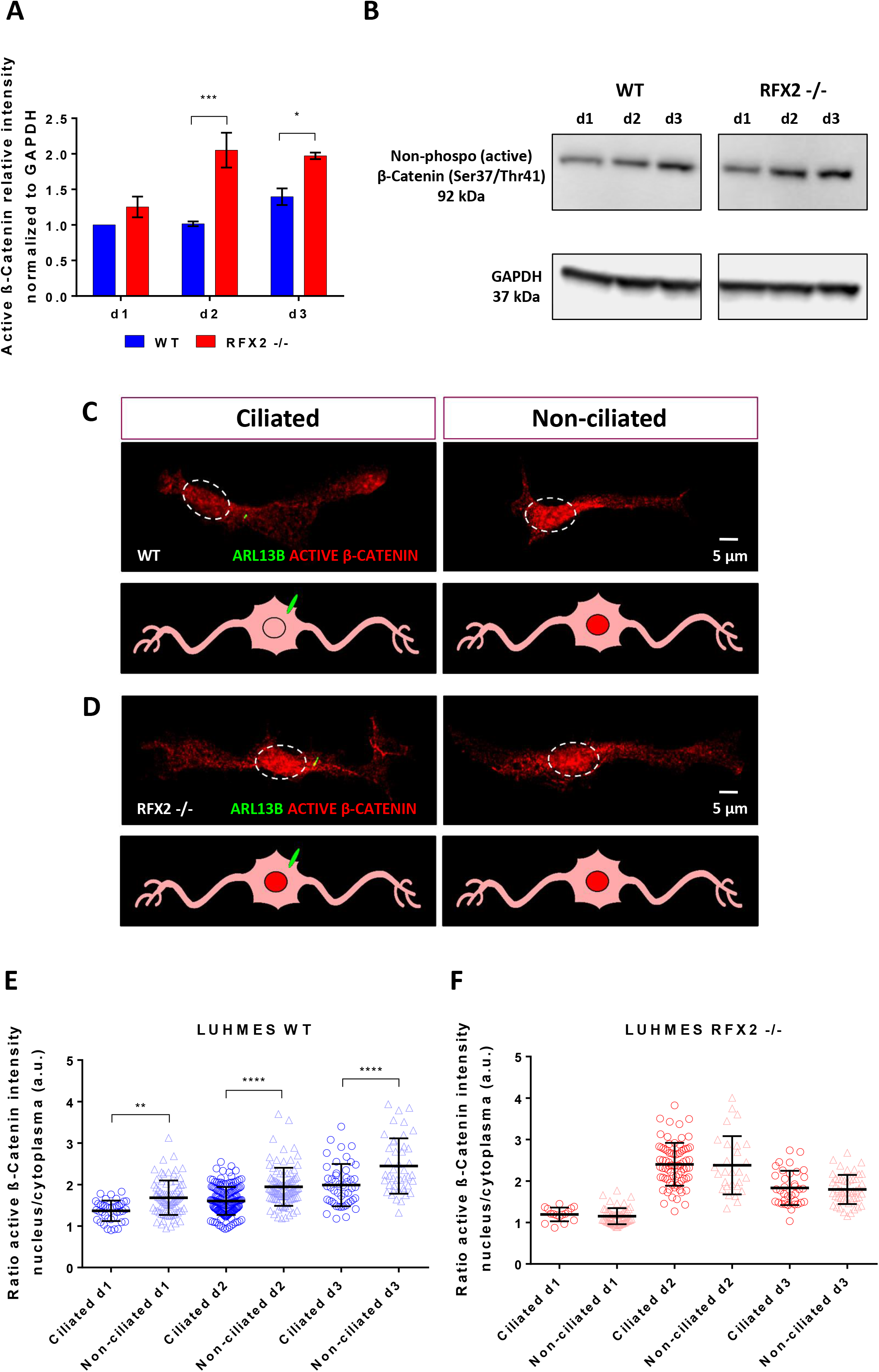
The intracellular localization of β-catenin, the major WNT signaling pathway transducer molecule, is regulated by the primary cilium. **(A-B)** The active (non-phosphorylated) form of β-catenin is significantly upregulated in an RFX2 -/- background as compared to WT neurons. This upregulation is particularly strong on d2 and d3 of neuron differentiation, the two time points with the highest percentage of ciliation (cf. Figure 1G). Quantification of expression values **(A)** was carried out from western blot analysis **(B)** relative to the housekeeping protein GAPDH (loading control). **(C-D)** Active β-catenin immunocytochemistry staining of WT and RFX2 -/- neurons highlights differences between genotypes in intracellular protein localization: note the more prominent nuclear localization in the RFX2 -/- background and in non-ciliated WT neurons in comparison to ciliated WT neurons. The ciliary marker is ARL13B (green). **(E-F)** Quantification of active β-catenin localization based on the ratio of immunofluorescence intensity between the cytoplasm and the nucleus on d1, d2 and d3 of LUHMES neuron differentiation. In WT, ciliated neurons more strongly promote β-catenin degradation in the cytoplasm, resulting in diminished translocation to the nucleus of the active form (lower ratio), as compared to non-ciliated neurons (higher ratio) **(E)**. In an RFX2 -/- background, ciliated and non-ciliated neurons show no differences in regulating β-catenin intracellular localization (equal ratios) **(F)**. Mean values are shown ± s.e.m (normalized to GAPDH) **(A)** and ± s.d. **(E-F)**. The results are from two independent experiments with a minimum of two technical replicates each. We conducted regular two-way ANOVA analyses (not repeated measures) with multiple comparisons (Bonferroni’s test) between groups. *p<0.05; **p<0.005; ***p<0.0005; ****p<0.0001.

To uncover the involvement of cilia in regulating the balance of cellular β-catenin localization, we performed immunocytochemistry to visualize where in differentiating neurons active β-catenin localizes. We compared fluorescence intensities (A.U.) detected in the nucleus and the surrounding cytoplasm and found that non-ciliated WT neurons were less efficient than ciliated neurons in inhibiting the nuclear activation of β-catenin (Figure 6C, 6E). While no β-catenin localization differences were detected in RFX2 -/- neurons between populations with altered cilia and non-ciliated populations (Figure 6D, 6F). Finally, consistent with the detection of bands on Western blots (Figure 6A, 6B), also immunocytochemistry revealed a stronger nuclear presence of active β-catenin in RFX2 -/- as compared to WT neurons (Figure 6D, 6F).

These findings suggest that cilia and ciliary signaling pathways do not regulate overall β-catenin protein levels but rather the subcellular turnover and localization of β-catenin. Furthermore, differentiation-delayed LUHMES RFX2 -/- neurons showed a higher activation of the β-catenin-mediated canonical WNT pathway, whereas LUHMES WT neurons appeared more efficient in switching to non-canonical WNT activation promoting neuron differentiation (Bengoa-Vergniory et al., 2014).

### WNT signaling is involved in the organization of the cyto-architecture

Neuron differentiation requires cytoskeletal rearrangements to form the outgrowth and branching of neurites (Clarke et al., 2020), and to generate synapses. Downstream effects of non-canonical WNT signaling can lead to modifications of the cyto-architectural organization (May-Simera and Kelley, 2012). To test whether the delayed differentiation observed in LUHMES RFX2 -/- neurons were due to a disrupted balance between canonical and non-canonical routes of the WNT signaling pathway with a consequent impairment of cytoskeletal structure, we treated neurons with a pathway inhibitor (Wnt-C59) or with a canonical pathway activator (Wnt3a) (Grumolato et al., 2010). The expression of WNT pathway output genes implicated in cyto-architectural functions (Luo et al., 2016; Khan et al., 2019) was subsequently assessed by qRT-PCR.

Treatment with a pathway inhibitor (Wnt-C59) revealed clear differences on how these WNT output genes responded: they were strongly downregulated in WT and only reached similarly basal expression levels in RFX2 -/- (Figure 7A). Aside from their structural functions, these genes are also widely known as axon markers (MAPT, TRIM46), non-canonical WNT pathway output genes (CAMK2A, DAAM1) and as candidate genes for neurodevelopmental disorders (CAMK2A, ANK2, DCX, SCN2A), highlighting their importance for cell functionality. Canonical WNT output genes were also tested following Wnt-C59 inhibition, but typically no significant differences in expression levels were detected (Figure 7C), further strengthening the importance for neuron differentiation of non-canonical WNT signaling (over the canonical route). Finally, the activation of the canonical WNT pathway with the activator Wnt3a showed a stronger upregulation of canonical target genes in WT, but not in the RFX2 -/- background (Figure 7B), where altered cilia may prove less efficient in detecting and transducing extracellular cues.

**Figure 7:**
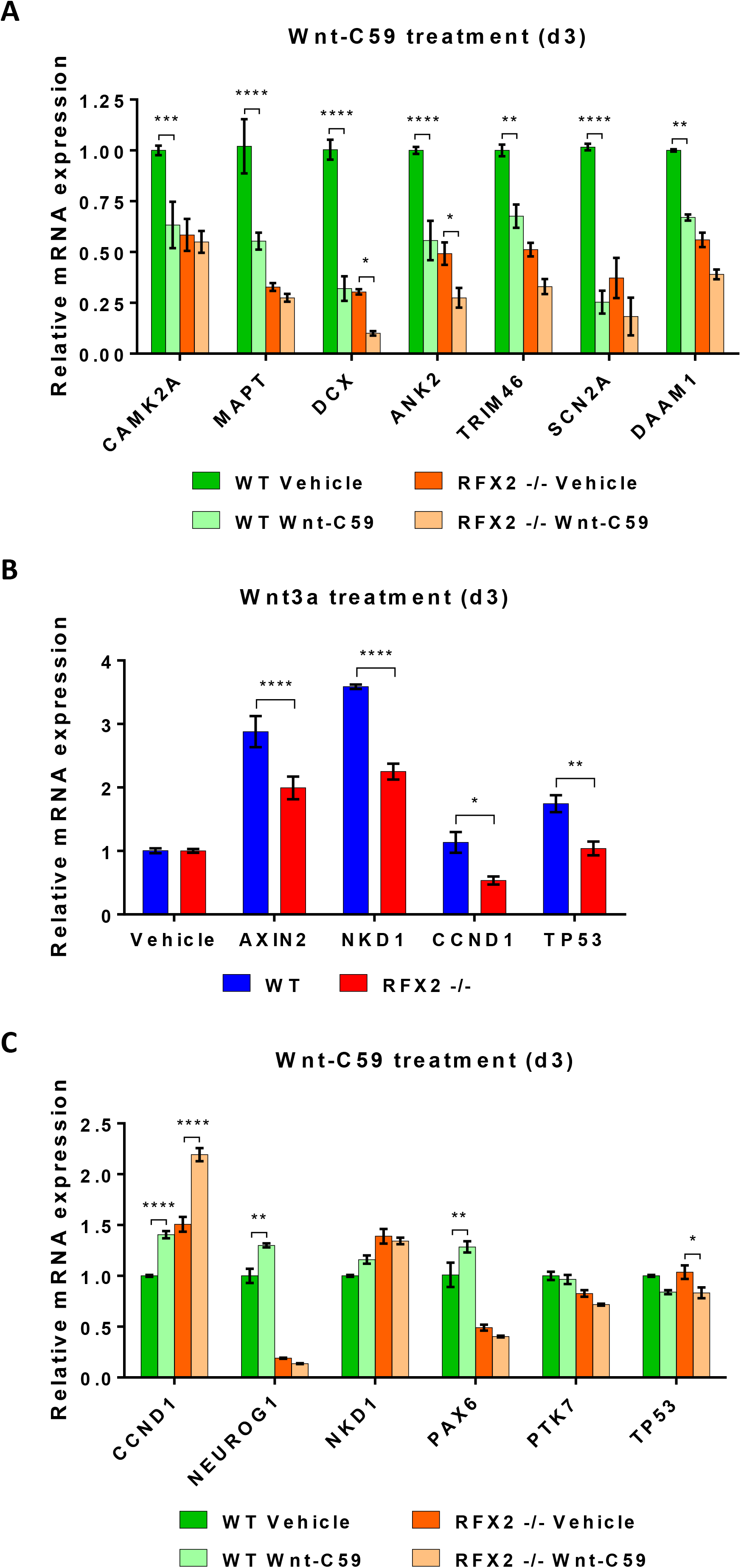
The expression of WNT signaling pathway output genes (with known or ascribed roles in cyto-architectural organization) is dependent on the primary cilium. qRT-PCR analysis of the expression of WNT signaling pathway output genes following pathway inhibition (antagonist Wnt-C59) or activation (agonist Wnt-3a) reveals clear gene expression differences in LUHMES WT neurons as compared to RFX2 -/- neurons with an altered primary cilium. **(A)** WT neurons treated with Wnt-C59 show strong downregulation of genes implicated in cyto-architectural organization, including the non-canonical WNT signaling pathway output genes CAMK2A and DAAM1: expression levels are very similar to the basal expression levels of untreated RFX2 -/- neurons (vehicle). **(B)** Treatment with a WNT activator (Wnt-3a) results in stronger upregulation of canonical output genes in WT neurons as compared to in the RFX2 -/- background. **(C)** qRT-PCR analysis of the expression of a set of canonical WNT signaling pathway output genes (Note: PTK7 is also as non-canonical target gene) following pathway inhibition (antagonist Wnt-C59): for this set of output genes WT neurons treated with Wnt-C59 do not show downregulation of gene expression and their expression levels are very similar to the basal expression levels of untreated (vehicle) RFX2 -/- neurons with an altered primary cilium (exceptions: neuron differentiation markers NEUROG1 and PAX6). These results point toward an involvement of the non-canonical WNT signaling pathway in neuron differentiation to promote cytoskeletal rearrangement and remodeling. Mean values are shown ± s.e.m (normalized to GAPDH; not shown). The results are from a minimum of three independent experiments. We conducted regular two-way ANOVA analyses (not repeated measures) with multiple comparisons (Bonferroni’s test) between groups. *p<0.05; **p<0.005; ***p<0.0005; ****p<0.0001.

These results describe molecularly how neuron differentiation changes and delays (axon outgrowth and branching), as observed in RFX2 -/- neurons, may arise in the absence of a fully functional signal transduction hub like the cilium: a deregulation of WNT signaling leads to impairment and defects in the cytoskeletal rearrangement capacities and consequently to impaired neuron differentiation.

## DISCUSSION

We investigated how cilia and ciliary signaling affect the development of the nervous system, using LUHMES, an established and validated human neuronal cell model (Scholz et al., 2011; Lauter et al., 2020), where we can differentiate proliferating neuronal precursor cells into fully mature, functional neurons within one week. We used wild-type (WT) and a CRISPR/Cas9 designed mutant for the ciliogenic transcription factor (TF) RFX2.

We could show that functional cilia and ciliary signaling are required to promote neuron maturation during an early differentiation time window: cilia promote axon outgrowth and regulate axon branching/arborization. These crucial aspects of differentiation were deregulated in neurons with mutated and altered cilia (RFX2 -/-). Microscopy and molecular data from STRT RNA-seq analyses demonstrated a delayed differentiation of RFX2 -/- neurons as compared to WT. When testing a highly relevant (ciliary) signaling pathway for neuron differentiation, WNT, we found that cilia play a key role in regulating the cellular turnover of β-catenin, the main mediator of WNT signaling, significantly decreasing its nuclear translocation and consequently, the canonical activation of the pathway.

In our assays the non-synchronized populations of post-mitotic, differentiating LUHMES neurons consisted of both ciliated and non-ciliated neurons. Our complete differentiation time course experiments from precursor to mature neuron revealed a clear pattern: ciliation steadily increased and peaked at d3 of differentiation (around 70% ciliation) before decreasing during later differentiation stages (d4-d6). Ciliation in post-mitotic, differentiating neurons is thus transient, suggesting a ciliary time window that appears to “open” *after* newborn neurons undergo the first centrosome-associated differentiation steps (polarization, growth cone development, initiation of neurites) (Meka et al., 2020; Meka et al., 2022) and to “close” (disassembly of cilia) *before* the emergence of large numbers of fully functional synapses as signaling hubs during the later neuron differentiation and maturation stages (Meka et al., 2020). This ciliary time window appears to be centered around differentiation stages 2 and 3 (breaking the bipolar symmetry of neurites and axon outgrowth). Importantly, we found that ciliation conferred an outgrowth advantage: ciliated neurons progressed more efficiently than non-ciliated neurons to stage 3 of differentiation. We note that our *in vitro* experimental setup did not allow for including the analysis of neuron migration, an important, and in parts concurrent, aspect of neuron differentiation and maturation.

To study whether and how ciliary alterations might affect neuron differentiation, we mutated the ciliogenic TF gene *RFX2*. In human, there are eight *RFX* genes (Sugiaman-Trapman et al., 2018). LUHMES cells and neurons prominently express several RFX family members with specific, but also potentially redundant roles (RFX1-3, RFX5, RFX7), where RFX5 and RFX7 have so far not been associated with ciliogenesis (Choksi et al., 2014). In LUHMES, *RFX2* is highly expressed at d0 and d1 of differentiation (our transcriptomics data) and RFX2 -/- cilia were not truncated but significantly longer than WT cilia. This suggests a potential role of the RFX2 TF in negatively regulating aspects of ciliogenesis at the onset of neuron differentiation, while RFX1 and RFX3, which are upregulated later, may be involved in regulating ciliogenesis after d1, when ciliary signaling promotes the differentiation process.

Our research aimed to uncover cellular and molecular mechanisms (when, where and how) by which cilia promote neuron development and differentiation. Our findings in WT that ciliated neurons more efficiently than non-ciliated neurons **(i)** reached differentiation stage 3 (break of symmetry, axon outgrowth) and **(ii)** generated significantly more branches on emerging axons, strengthens the concept that cilia are involved in critical, axon-associated aspects of neuron differentiation. By contrast, altered cilia of RFX2 -/- neurons did not provide those differentiation advantages over non-ciliated neurons: fewer neurons were able to reach stage 3 of differentiation and there were no significant branching differences between the emerging axons of ciliated and non-ciliated neurons. Interestingly and differently from WT, RFX2 -/- axons from non-ciliated neurons were as highly branched as neurons with altered cilia, again implicating RFX2 in a negative regulatory role regarding axon branching, which in turn may later facilitate axon pruning. The specificity of the ciliary impact on neuron (axon) differentiation is further underscored by the fact that populations of WT and RFX2 -/- neuronal precursor cells and neurons did not display any relevant differences in proliferation rates nor in neuron types (bipolar neurons dominated in populations of both genotypes).

We complemented our cellular observations with analyzing the STRT RNA-seq transcriptomes of both genotypes in time course experiments. RFX2 -/- neurons were delayed in differentiation as compared to WT neurons at the same time point. Ciliary alterations (like in RFX2 -/-) therefore seem to produce a relatively mild phenotype (as it frequently is the case in ciliopathies), where neurons differentiate like in WT, but with a different timing, which may disrupt critical spatiotemporal aspects of brain formation. Gene expression profiles from transcriptomics were grouped into three macro-clusters, whereby we particularly focused on Cluster 3 because the GO analysis of sets of downregulated genes in RFX2 -/- neurons showed strong enrichments of functional characteristics related to the nervous system, neuron and axon development, explaining the RFX2 -/- delay in neuron differentiation at the molecular level. Moreover, sequence motifs associated with downregulated genes were strongly enriched for binding sites of TFs prominently involved in WNT signaling, a cilia-associated pathway well-known for regulating neuron development (Inestrosa and Varela-Nallar, 2015; Arredondo et al., 2020).

Different types of small molecules are available to modulate WNT signaling at different levels of the pathway (Tran and Zheng, 2017). LUHMES neurons are an *in vitro* model where the culture conditions do not fully reflect an *in vivo* extracellular environment with exogenous WNT ligands. As such, the WNT pathway in LUHMES is mainly regulated by endogenous WNT signals produced in the cell. Furthermore, both the canonical and non-canonical pathway routes (Wheway et al., 2018) may simultaneously be operative. Therefore, we treated differentiating LUHMES neurons with the Wnt-C59 antagonist (Motono et al., 2016) to inhibit the pathway in full, preventing secretion and subsequent activation of all endogenous WNT signals, regardless of their pathway affinity. In WT neurons this resulted in a significant reduction of axon outgrowth (stage 3 neurons) irrespective of the presence or absence of cilia, and increased and deregulated axon branching, a phenotype also observed in the RFX2 -/- background. RFX2 -/- neurons treated with Wnt-C59 or vehicle (control) showed no differences between ciliated and non-ciliated neurons in both conditions, with a decreased percentage of stage 3 neurons as compared to WT and a similar upregulation of axon branching in non-ciliated neurons in both experimental conditions. These results clearly demonstrate that WNT signaling affects neuron differentiation and axon outgrowth, potentially coordinating the formation of necessary *versus* eventually unnecessary branches, a key aspect of subsequent neuron interconnectivity and circuitry.

To elucidate where and how cilia regulate WNT signaling, we determined the cellular localization of the active form of β-catenin, noticing in WT a significant reduction of the ratio nucleus/cytoplasm in the presence of cilia, while in RFX2 -/- with altered cilia and thus, apparent modulation deficits, ciliated and non-ciliated neurons displayed similarly high ratios nucleus/cytoplasm. These results reveal that cilia are important for controlling (and reducing) the nuclear translocation of β-catenin rather than mediating its cytoplasmic degradation. Since β-catenin is the main mediator of canonical WNT signaling, this indicates that in (LUHMES) neurons it is functional cilia that reduce the activation of canonical WNT signaling, to favor the non-canonical route for promoting the rearrangement of cytoskeletal structures required to develop and grow neurites and branches during early neuron differentiation (Bengoa-Vergniory et al., 2014). Further, inhibition with Wnt-C59 also led to a strong downregulation of key genes involved in cytoskeletal remodeling (Godoy et al., 2021), many of which were similarly downregulated to basal expression in RFX2 -/-. In addition, by exposing LUHMES neurons to an exogenous canonical WNT signaling pathway agonist (Wnt3a) (Janda et al., 2017) we found that WT neurons (with WT cilia) were better equipped than RFX2 -/- neurons (with altered cilia) to upregulate the expression of canonical WNT signaling target genes. In summary, ciliary regulation of WNT signaling and its output proves essential to drive neuron differentiation by translating signaling events into anatomical changes necessary to shape the emerging axon and its branching pattern. Testing further combinations of pathway activator/inhibitor molecules will in the future be needed to dissect and resolve the when and where of ciliary regulation of WNT signaling output more precisely.

Based on the pattern of ciliation in LUHMES neurons (steadily increasing and peaking around d3 of differentiation before decreasing again; particularly prominent at differentiation stages 2 and 3), we propose the following working model, integrating previous and our current results. The centrosome functioning as MTOC leads the initiation of neuron differentiation, driving precursor cell polarization and determining the axon formation site (Meka et al., 2020; Meka et al., 2022). Ciliogenesis ensues and thereby informative ciliary signaling becomes crucially important during an early differentiation time window to promote axon outgrowth (breaking the bipolar symmetry during differentiation stages 2 and 3), axon branching and arborization. Later, during differentiation stages 4 and 5, ciliation declines and the establishment of many functional synapses (across a now much larger anatomical space) take over relevant signaling tasks for completing neuron differentiation and maturation (Liu et al., 2021). We note that our working model does not consider neuron migration, a developmental process not tested, but which *in vivo* partially overlaps with the anatomical differentiation processes described here (neuron migration has also been shown to be subject to ciliary regulation – see below). Mechanistically, the ciliary impact on neuron differentiation is executed by tipping the balance from canonical to non-canonical WNT signaling, decreasing the nuclear translocation of β-catenin and, therefore, initiating cytoskeletal rearrangements to promote and sustain neuron differentiation.

Our previous (Lauter et al., 2020) and current work in human LUHMES neurons on the ciliary impact on axon outgrowth and branching compares well with complementary work in other systems. Cilia and ciliary signaling were found to promote axonal growth cone architecture and axon tract development in mouse through the regulation of the PI3K/AKT signaling pathway (Guo et al. 2019). Cilia and ciliary signaling proved essential for dendrite outgrowth and refinement as well as for the initiation of synapse formation (Kumamoto et al., 2012; Guadiana et al., 2013). Stoufflet and colleagues (2020) described cilia as the crucial organelle for driving *in vivo* saltatory migration of neurons in the mouse brain through the cAMP/PKA pathway, being cyclically extended in the leading process (future dendrite) of migration. Whereby, disruption of neuronal ciliogenesis can lead to defective neuron migration and inactivation of non-canonical WNT signaling in association with aberrant cytoskeletal remodeling (May-Simera and Kelley, 2012; Park et al., 2019).

Our work expands on previous studies as we provide in a human neuronal cell model, LUHMES, detailed and complete, cellular and molecular time course analyses of neuron development from proliferating neuronal precursor cell to differentiating and fully differentiated, mature neuron. In LUHMES it is possible to precisely analyze the function of cilia and ciliary signaling at every single step of neuron differentiation. Moreover, we have elucidated a molecular mechanism underlying the ciliary regulation of cytoskeletal rearrangements necessary to promote neuron differentiation through the activation of non-canonical WNT signaling. It is highly likely that WNT signaling cross-talks with other relevant ciliary signaling pathways like PI3K/AKT and cAMP/PKA (Mercado-Gómez et al., 2008; Zhang et al., 2013). Thus, to fully understand the ciliary regulation of neuron differentiation, investigating other (interacting) ciliary signaling pathways, on top of our discovery of WNT signaling, will likely be required.

Dysregulation of ciliary activity has increasingly been found to be implicated in neurodevelopmental conditions and disorders (Valente et al., 2014; Reiter and Leroux, 2017; Lauter et al., 2018). We hypothesize an early differentiation time window where cilia and ciliary signaling drive the anatomical transition from proliferating precursor to differentiated neuron, by promoting neuron migration and axon development needed for proper brain formation. We propose that ciliary malfunctions result in an inability to properly translate signaling events into anatomical changes, which can lead to impaired neuron migration to eventual brain destinations or to anatomical deficits, eventually causing localized circuitry defects and increasing the risk of the onset of certain neurodevelopmental conditions and disorders like autism, schizophrenia and dyslexia (Massinen et al., 2011; Lauter et al., 2018; Harris et al., 2021).

### LIMITATIONS OF OUR STUDY

LUHMES represents a simplified *in vitro* system comprised of a single type of neuron, lacking the support of and interactions with glial and endothelial cells that occur *in vivo*. In regular cell culture setups LUHMES affords very limited possibilities for investigating cell and neuron migration, a very important “partner” in neuron differentiation. While future 3D brain organoid setups may be needed to fully understand and explore the ciliary contributions to brain formation, several *in vitro* approaches have already been established as valuable tools to study neuron migration at the cellular and molecular level and these can be adjusted for being applied in LUHMES neuron cultures as well (Azzarelli et al., 2017). However, a simplified system may represent an advantage when it comes to isolate, clearly separate, and analyze specific neuron differentiation steps. LUHMES neurons make possible to analyze the complete neuron differentiation process in unprecedented detail at every single step. A necessary system upgrade will be the ability to continuously follow individual fluorescently marked cells or neurons over time by using live cell imaging approaches (transient or permanent transgenesis) to uncover in minute detail the flow (rather than discrete steps) of cilia-promoted neuron differentiation. Live visualization of individual neurons will make possible the accurate molecular analyses by single cell RNA-sequencing through the ability to selectively pick ciliated and non-ciliated cells and neurons from the same culture. LUHMES is a hard-to-transfect cell line, though reasonably good results have been obtained using novel electroporation protocols (Shah et al., 2016; Calamini et al., 2021), making LUHMES suitable for genome editing by CRISPR/Cas9 at a larger scale.

## MATERIALS AND METHODS

### Cell culture and growth conditions

<u>Lu</u>nd <u>h</u>uman <u>mes</u>encephalic (LUHMES) cells, a v-myc immortalized neuronal precursor cell line, was obtained from the ATCC (https://www.atcc.org/products/crl-2927). LUHMES cells were cultured in a standard incubator (37°C, 5% CO_2_) as previously described (Scholz et al., 2011; Lauter et al., 2020; Coschiera et al., 2023). LUHMES cells were grown in DMEM/F-12 Ham growth medium (Sigma-Aldrich D6421) supplemented with L-glutamine solution (Sigma-Aldrich G7513; 2.5 mM), N-2 supplement (Gibco 17502-048; 1×) and human heat stable basic Fibroblast Growth Factor (bFGF) (Thermo Fisher Scientific PHG0369; 20 ng/ml) in vessels pre-coated with poly-L-ornithine hydrobromide (Sigma-Aldrich P3655, 50 μg/ml) and fibronectin from human plasma (Sigma-Aldrich F1056; 1 μg/ml). For some experiments, a higher concentration of ornithine (100 μg/ml), fibronectin (10 μg/ml) and bFGF (40 ng/ml) was used to improve cell adhesion and propagation. To differentiate LUHMES neuronal precursor cells into post-mitotic neurons, bFGF in the growth medium was replaced with tetracycline hydrochloride (Sigma-Aldrich T7660; 1 μg/ml) to terminate v-myc transgene expression.

### Immunocytochemistry and fluorescence intensity quantification

Immunocytochemistry was performed as previously described (Lauter et al., 2020; Coschiera et al., 2023). 7.5×10^4^ LUHMES cells were seeded 24h prior to differentiation into pre-coated 35 mm dishes (Corning 353001) containing round borosilicate cover glasses (VWR 631-0150P), pre-treated with hydrochloric acid fuming 37%, rinsed with dH_2_O and absolute ethanol and finally stored in 70% ethanol. Cells were fixed in 2% paraformaldehyde (PFA) solution (Invitrogen FB002) for 5 min in the incubator and permeabilized with 0.2% Triton X-100 in PBS for 12 min at room temperature (RT). The glass cover slips were then incubated with blocking buffer (2% bovine serum albumin and 0.1% Tween 20 in PBS) for 30 min at RT, prior to primary antibody incubation overnight at 4°C and secondary antibody incubation for 1h at RT. Cells on glass cover slips were rinsed twice in PBS before being mounted on microscope slides (Thermo Fisher Scientific 15457544) using 4 μl of ProLong glass antifade mountant (Invitrogen P36982). Samples were then left in the dark on a flat surface for 24h at RT to allow them to cure and reach the optimal refraction index.

All primary and secondary antibodies were diluted in blocking buffer and used under the following working conditions. Neuron differentiation markers were chicken anti-TAU (MAPT) (Abcam ab75714; 1:100), rabbit anti-TRIM46 (Human Protein Atlas HPA030389; 1:100), mouse anti-TUBB3 (TUJ1) (BioLegend 801213; 1:3000) and rabbit anti-PSD95 (Cell Signaling 2507; 1:100). Centrosome (centriole, basal body) and cilium markers were, respectively, mouse anti-PCNT (Abcam ab28144; 1:250) and rabbit anti-ARL13B (Proteintech 17711-1-AP; 1:10000). Mouse anti-active-β-catenin (Sigma-Aldrich 05-665; 1:500) was used to detect the main cellular mediator of WNT signaling. The intensity of active-β-catenin fluorescence was measured as mean value in arbitrary units (A.U.) and quantified using Fiji software (Schindelin et al., 2012). Secondary antibodies were conjugated with Alexa Fluor dyes 488, 555, 647 (Invitrogen) and used at a 1:600 dilution. Alexa Fluor 647 Phalloidin staining (Invitrogen A30107; 1×) was coupled with secondary antibody incubation to visualize F-actin. Nuclei were stained for a few seconds with the Hoechst dye 33342 (Invitrogen H3570; 50 μg/ml in PBS).

### Microscopy and image acquisition

Microscopy work was performed at the Live Cell Imaging Core facility/Nikon Center of Excellence, at the Karolinska Institute (https://ki.se/en/bionut/welcome-to-the-lci-core-facility). For immunocytochemistry we used a Nikon Ti-E inverted spinning disk CREST v3 confocal microscope with a fully motorized piezo stage for 3D z-stacking and a Plan Apo λ 60× NA 1.40 oil immersion objective, matched to the refractive index of the sample mounting medium and cover glasses. Images were acquired with a Kinetix sCMOS camera (>95% quantum efficiency, 6.5 μm pixel size, 2720×2720 pixels field of view).

Measurements of differentiating axons: Neurites were marked with anti-TAU or anti-TRIM46 as axon markers. The length and diameter of neurites were measured using the line tool of the Fiji software (Schindelin et al., 2012). Neurite length was measured as the trajectory starting at the center point of the nucleus (stained with Hoechst dye 33342) and reaching the tip of the neurite. Only neurons with the main neurite (emerging axon) being at least 1.5 times longer than the opposite neurite were included in the differentiation stage 3 group of neurons: after the break of bipolar symmetry.

Cilia length measurements: Cilia in LUHMES emanate from the neuronal cell body (Lauter et al., 2020). We marked ciliary basal bodies with anti-PCNT and the entire ciliary shaft with anti-ARL13B. We used a microscopy z-stack imaging setup, in which cilia were freely “floating” in the mounting medium: cilia were physically not obstructed in any of the XYZ axes by neither the glass slide nor by the cover glass (Coschiera et al., 2023). The length of individual cilia identified by anti-ARL13B fluorescence was then measured base-to-tip using the line tool of the Fiji software (Schindelin et al., 2012).

### Drug treatment assays (Wnt-C59, Wnt-3a)

3.5×10^5^ LUHMES cells were seeded in a pre-coated 6-wells plate (Corning 353046) for growth overnight before switching to the differentiation culture medium (d0). Wnt-C59 (Tocris 5148), an inhibitor of WNT signaling (Motono et al., 2016), was used at 2 µM for a 72h incubation step (from d0 until d3 of neuron differentiation). The WNT signaling pathway activator Wnt-3a (Kaur et al., 2013) (R&D Systems 5036-WN-010) was instead supplied at 200 ng/ml for 24h (on d1), before adding another 200 ng/ml on d2, followed by 24h incubation (until d3, 48h total incubation). As control, LUHMES cells were cultured following the same differentiation and drug incubation procedures, whereby only 0.2% DMSO and PBS (0.1% BSA) were added, as these were used as vehicle for Wnt-C59 and Wnt-3a, respectively.

### CRISPR/Cas9 mutation of the RFX2 gene in LUHMES cells

We generated an RFX2 (-/-) mutant LUHMES cell line by using CRISPR/Cas9-mediated non-homologous end joining. To minimize off-target effects the double-nicking strategy with the Cas9 D10A nickase mutant was used in combination with paired guide RNAs (Ran et al., 2013) (for sgRNA sequences see Supplementary Table S5). We double-checked for potential off-target effects by using software (https://cctop.cos.uni-heidelberg.de:8043; Stemmer et al., 2015) that determines and analyzes the predicted number of sequence mismatches and their distance from the PAM sequence that would need to be tolerated by the Cas9 enzyme. The only gene with 0 mismatches with the designed sgRNAs was RFX2. All the other genes listed as off-target candidates had at least 4 mismatches, including in the PAM sequence “core region”. Annealed oligonucleotides covering the 20 nt guide sequences and an NGG PAM site were then cloned into the BbsI restriction site of the plasmid pSpCas9n(BB)-2A-GFP (pX461; Addgene plasmid #48140; gift from Feng Zhang). The resulting plasmid constructs were verified by Sanger-sequencing prior to use.

Approximately 1×10^6^ LUHMES cells were seeded into coated 10 cm petri dishes in complete medium (Sigma-Aldrich D6421). Transfection was carried out the following day at about 40% confluency. 7.5 μg of each sgRNA containing plasmid (15 μg in total) were transfected together with 24 μl lipofectamine (Invitrogen 18324012) in 5 ml fresh complete medium overnight. The following day fresh complete medium was re-supplied and cells were allowed to recover for 48-72h. For cell-enrichment, transfected LUHMES cells were FACS-sorted and GFP expressing cells were collected for further culture. Prior to sorting, cells were trypsinized and singled-out in PBS, followed by a passage through a wet cell strainer (mesh size 40 μm). For clone isolation approximately 1000 FACS-sorted GFP-positive cells were seeded per 10 cm dish in 8 ml complete medium and cultured for 10 days with one medium change. On day 10 single cell-derived colonies were picked up using a 100 μl pipette tip and transferred into a 96-well plate containing complete medium. Clones were sub-cultured continuously until the plate wells were fully covered. At this point replicas of isolated clones were generated and sub-cultured in a 24-well plate format. One replica was used for further culturing, while the other one was used to screen for mutations in the RFX2 gene.

For screening, cells were trypsinized and pelleted by centrifugation. Extraction of template genomic DNA was performed by using QuickExtract (Epicentre QE0905T). 2 μl of genomic DNA extract solution was used to PCR amplify the genomic locus of interest (RFX2) with an intronic primer pair. PCR-products were subsequently Sanger-sequenced for the identification of mutations in the RFX2 gene. Location and sequence details of the CRISPR/Cas9-generated mutations in the RFX2 gene are shown in Supplementary Figure S3.

### Proliferation rate assay in different culture conditions

To assess cell proliferation rates in different conditions (proliferation versus differentiation), both wild type and RFX2 -/- LUHMES cells were seeded in parallel in pre-coated 6-wells plates (Corning 353046) for growth overnight in regular proliferation medium until the next day (time point = 0h). To assess growth in regular proliferation conditions, cells were grown for an additional 72h until they became fully confluent. To assess growth under neuron differentiation conditions, proliferation medium was changed to differentiation medium at 24h, and cells were kept in culture for 96h in total. Images at different time points of culture were acquired using a bright-field microscope (Carl Zeiss, Axiovert 200M) and then transferred into Fiji software (Schindelin et al., 2012) for manual cell counting.

### Western blot and protein expression quantification (RFX2, active β-catenin)

Wild type and RFX2 -/- LUHMES cells were cultured in a T75 flask until reaching 80% confluency. Cells were then washed twice with ice cold PBS and mechanically detached using a cell scraper (Sarstedt 83.3951) in RIPA lysis buffer (Millipore 20-188; 1×) 1% SDS solution after adding protease inhibitor (Roche 4693116001; 1×) and phosphatase inhibitor cocktails (Roche 4906845001; 1×). Cell samples were disrupted by vortexing, using a syringe, and clarified centrifugation at 14000×g for 15 min at 4°C, followed by supernatant collection in a new Eppendorf tube. Protein concentration was determined using a BCA assay kit (Thermo Fisher Scientific 23227) and a microplate reader (Tecan M200 Infinite Pro). Protein samples were then mixed with LDS loading buffer (Invitrogen NP0007; 1×) and reducing agent (Invitrogen NP0009; 1×) and denatured at 70°C for 10 min. 20 μg of protein sample and a reference ladder (Bio-Rad 1610374) were loaded in NuPage 4%-12%, Bis-Tris gel (Invitrogen NP0321PK2) and electrophoresis was carried out in MOPS SDS buffer (Invitrogen NP0001; 1×) at 200 V for 1h. At the end of the gel run, transfer buffer was prepared (for 1 L: 14.4 g glycin, 3.03 g Tris Base, 200 ml methanol, up to volume with dH_2_O), the nitrocellulose membrane was activated in methanol for 5 min prior to the assembly of the transfer sandwich; the protein transfer was run at 100 V for 1h at 4°C. The membrane was retrieved from the sandwich and washed with PBS-T (0.1% Tween 20) for 5 min, before being blocked with 5% non-fat dry milk in PBS-T for 1h and incubation with the primary antibody overnight at 4°C on a shaker. The membrane was then washed three times in PBS-T and incubated with the secondary antibody for 1h at RT on a shaker.

All primary and secondary antibodies were diluted in 5% non-fat dry milk in PBS-T. Two primary rabbit anti-RFX2 antibodies were successfully tested and used at a 1:1000 dilution (Atlas Antibodies HPA048969 – Figure 2B; Thermo Fisher Scientific PA5-61850 – data not shown). Mouse anti-active-β-catenin was used at 1:2000 (Sigma-Aldrich 05-665 – Figure 6B-D). Rabbit anti-GAPDH was used as housekeeping reference protein antibody (Invitrogen PA1-987; 1:1000). The secondary antibodies were horseradish peroxidase (HPR)-conjugated (Sigma-Aldrich A0545, anti-rabbit 1:10000; GE Healthcare NA931V, anti-mouse 1:5000). Protein expression was detected after 3 min incubation with a chemoluminescent substrate (GE Healthcare RPN2106 for GAPDH; Thermo Fisher Scientific 80196 for active-β-catenin; Thermo Fisher Scientific 34095 for RFX2). Images were taken with the ChemiDoc Touch Imaging System (Bio-Rad) and bands were quantified with the Fiji software (Schindelin et al., 2012).

### Isolation of total RNA

Total RNA was extracted from wild type and RFX2 -/- LUHMES cells and column-purified using an RNeasy mini kit (Qiagen 74104) following the manufacturer’s protocol. RNA integrity was checked in 1% agarose gel electrophoresis (Sigma-Aldrich A9539) and concentration and purity were assessed with a Nanodrop ND-100 (Thermo Fisher Scientific). For library preparation and transcriptome analysis, an additional DNase step (Qiagen 79254) was introduced during RNA isolation. RNA integrity values were evaluated using an Agilent Tech 2200 Tape Station (RIN value >8; KI Bioinformatics and Expression Analysis core facility; https://ki.se/en/bionut/bea-core-facility). RNA concentration was determined with a Qubit 3.0 Fluorometer (Thermo Fisher Scientific). The samples were then diluted to 10 ng/μl (±0.5 ng/µl) in nuclease-free water before proceeding with the RNA-seq library preparation.

### cDNA synthesis and quantitative real-time PCR (qRT-PCR)

1 μg of total RNA was used to synthesize cDNA using a 1:1 mix of oligo (dT) and random hexamer primers of a RevertAid H Minus First Strand cDNA Synthesis Kit (Thermo Fisher Scientific K1631). The cDNA product was further diluted 1:4 in nuclease-free water and 2 μl were used in a 15 μl total reaction for qRT-PCR (7500 Fast Real-Time PCR instrument; Applied Biosystems), using SYBR green as nucleic acid stain (Roche 4913850001). To specifically quantify mRNA expression, both forward and reverse primers (Supplementary Table S5) were designed on consecutive exons and, during the qRT-PCR run, a melting curve analysis was performed to ensure specific amplification of the respective gene of interest. Relative mRNA levels were evaluated using the 2^−ΔΔ^ ^Ct^ method and GAPDH was used as housekeeping gene to normalize the values (Livak and Schmittgen, 2001).

### STRT RNA-seq library preparation and sequencing

We used 20 ng of RNA to generate two 48-plex RNA-seq libraries employing a modified STRT method with unique molecular identifiers (UMIs) (Islam et al., 2011, 2014). Briefly, RNA samples were placed in a 48-well plate, and a universal primer, template-switching oligonucleotides and a well-specific 6-bp barcode sequence (for sample identification) were added to each well of the plate (Krjutškov et al., 2016). The synthesized cDNAs from these samples were then pooled into one library and amplified by single-primer PCR using a universal primer sequence. STRT library sequencing was carried out with an Illumina NextSeq 500 System, High Output (75 cycles), at the Biomedicum Functional Genomics Unit (FuGU), University of Helsinki, Finland (https://www.helsinki.fi/en/infrastructures/genome-analysis/infrastructures/biomedicum-functional-genomics-unit).

### STRT RNA-seq data preprocessing

The raw STRT sequencing data in the Illumina Base Call Format were demultiplexed and processed using the STRT2 pipeline (Ezer et al., 2021) (https://github.com/my0916/STRT2). Reads were aligned to the human reference genome hg19, the human ribosomal DNA unit (GenBank: U13369) and ERCC spike-ins (SRM 2374) using GENCODE (v28) transcript annotation by HISAT2 (v2.2.1) (Kim et al., 2019) with the option ‘–dta’. For gene-based analyses, uniquely mapped reads within the 5’ UTR or 500 bp upstream of protein-coding genes were counted using Subread feature Counts (v2.0.0). For transcript far 5′ end (TFE) based analyses, the mapped reads were assembled using StringTie (v2.1.4) (Pertea et al., 2015), and mapped reads within the first exons of the assembled transcripts were counted as previously described (Töhönen et al., 2015). FASTQ files after exclusion of duplicated reads were deposited in the ArrayExpress database at EMBL-EBI (https://www.ebi.ac.uk/arrayexpress): accession number E-MTAB-11546. Numbers of total and mapped reads for each sample are summarized in Supplementary Table S1.

### Downstream expression analysis

Among the 80 samples from wild type and RFX2 -/- LUHMES cells and neurons, two samples were excluded as outliers due to extremely low numbers of total read counts. After filtering out lowly expressed genes and ERCC spike-ins, differential expression analysis was performed with the R (v4.0.3) package DESeq2 (v1.30.1) (Love et al., 2014). Genes or TFEs with an adjusted P-value of less than 0.05 were considered as significantly differentially expressed. Gene ontology (GO) term enrichment analysis was performed using the R package enrichR (v3.0) (Kuleshov et al., 2016). Motif enrichment on differentially expressed TFEs was analyzed using the command findMotifsGenome.pl from HOMER (v4.11) (Heinz et al., 2010) with the option ‘-size -300,100 -mask’, using all detected TFEs as background. Principal component analysis (PCA) was carried out using the 500 genes with the highest variance across all 78 samples with the DESeq2 PCAplot function. Time-course clustering was performed using the R package maSigPro (v1.62.0) (Nueda et al., 2014). Gene set enrichment analysis (GSEA) was performed with the R package fgsea (Korotkevich et al., 2021), where genes pre-ranked based on their P-values and fold changes were compared with the gene set of 686 ciliary genes from the SYSCILIA gold standard (SCGSv2) (Vasquez et al., 2021). Integrative analysis with the fetal midbrain single-cell RNA-seq data (GSE76381) (La Manno et al., 2016) was carried out with the R package SingleR (v1.4.1) (Aran et al., 2019), using the log2-normalized expression levels of the LUHMES STRT RNA-seq data as the reference data set.

## ACKNOWLEDGEMENTS

Parts of this study were performed at the Karolinska Institute Live Cell Imaging facility supported by grants from the Knut and Alice Wallenberg Foundation, the Swedish Research Council, the Center for Innovative Medicine, and the Jonasson Center at the Swedish Royal Institute of Technology. We acknowledge BEA, the Bioinformatics and Expression Analysis core facility, which is supported by the board of research at the Karolinska Institute and the research committee at the Karolinska Hospital. The Biomedicum Functional Genomics Unit at the University of Helsinki provided sequencing services. All computation for this work was performed on resources provided by the Swedish National Infrastructure for Computing (SNIC) at the Uppsala Multidisciplinary Center for Advanced Computational Science (UPPMAX) partially funded by the Swedish Research Council through grant agreement No. 2018-05973.

## AUTHOR CONTRIBUTIONS

Project conceptualization and planning: AC, MY, GL, JK, PS

Experimentation and methodology: AC, MY, GL, SE

Data analysis: AC, MY, GL, SE, JK, PS

Critical resources and reagents: AC, MY, GL, SE, JK, PS

Writing and illustrations – draft: AC, MY, PS

Writing and illustrations – editing and review: AC, MY, GL, SE, JK, PS

Supervision and project management: JK, PS

Funding acquisition: JK, PS

